# Deep Coverage and Extended Sequence Reads Obtained with a Single Archaeal Protease Expedite *de novo* Protein Sequencing by Mass Spectrometry

**DOI:** 10.1101/2025.05.26.656138

**Authors:** Tatiana M. Shamorkina, Laura Pérez Pañeda, Tereza Kadavá, Douwe Schulte, Patrick Pribil, Sibylle Heidelberger, Allison Michele Narlock-Brand, Steven M. Yannone, Joost Snijder, Albert J. R. Heck

## Abstract

The ability to sequence proteins without reliance on a genomic template defines a critical frontier in modern proteomics. This approach, known as *de novo* protein sequencing, is essential for applications such as antibody sequencing, microbiome proteomics, and antigen discovery, which require accurate reconstruction of peptide and protein sequences. While trypsin remains the gold-standard protease in proteomics, its restricted cleavage specificity limits peptide diversity. This constraint is especially problematic in antibody sequencing, where the functionally critical regions often lack canonical tryptic sites. As a result, exclusively trypsin-based approaches yield sparse reads, leading to sequence gaps. Multi-protease and hybrid-fragmentation strategies can improve the sequence coverage, but they add complexity, compromise scalability and reproducibility.

Here, we explore two HyperThermoacidic Archaeal (HTA)-proteases as single-enzyme solutions for *de novo* antibody sequencing. Each HTA-protease generated about five times more unique peptide reads than trypsin or chymotrypsin, providing high redundancy across all CDRs. Combined with EAciD fragmentation on a ZenoTOF 7600 system, this approach enabled complete, unambiguous antibody sequencing. *De novo* analysis using PEAKS/DeepNovo and Stitch showed up to fourfold higher alignment scores and reduced the sequence errors within the HTA-generated data. Additionally, the HTA-EAciD approach offers short digestion times, eliminates extensive cleanup, and enables analysis in a single LC-MS/MS run. This streamlined, single-protease approach delivers therefore performance comparable to multi-enzyme workflows, offering a scalable and efficient strategy for *de novo* protein sequencing across diverse applications.

## Introduction

Mass spectrometry (MS)-based proteomics has matured immensely, tackling challenging areas such as single cell proteomics, spatial proteomics, clinical proteomics and proteoform analysis (1, 2). However, most of the above-mentioned approaches are strongly dependent on reference databases for protein identification. A next frontier in MS-based proteomics is template-free protein sequencing which circumvents the need for a genomic database. This approach, known as *de novo* sequencing, provides sequence information directly from MS data and it is of both fundamental interest to the MS community and of practical importance in scenarios where genomic references are unavailable. Key applications of *de novo* sequencing include antigen discovery, antibody sequencing, and microbiome proteomics, all of which require accurate reconstruction of peptide and protein sequences (3, 4).

In standard peptide-centric proteomics, proteins are typically identified and quantified based on a few peptides generated by enzymatic digestion, typically using trypsin (5, 6). Trypsin cleaves strictly C-terminal to lysine and arginine residues, commonly generating ∼10 amino acid peptides in a narrow 700–1,500 Da mass range (5–9). Tryptic peptides are well-suited for most bottom-up (BU) analysis due to their favorable ionization and fragmentation properties when using collision induced dissociation (CID) (6–8, 10). However, achieving complete protein sequence coverage with tryptic peptides is rarely possible (6, 7). This limitation is particularly restrictive for the *de novo* workflows, especially in complex applications like *de novo* antibody sequencing (11–16).

The human antibody repertoire is immense, comprising proteins with shared architecture and structure, albeit each carrying a unique sequence. It has been estimated that the human adaptive immune system can generate over 10^15^ antibodies of distinct sequence (16–19), far exceeding the typically quoted ∼20,000 protein-coding genes (20, 21). A major class of human antibodies, IgGs, consists of two identical heavy (Hc) and two identical light chains (Lc) (17, 21). Each heavy-light chain pair forms one Fab (fragment antigen-binding) region, responsible for antigen binding. Within each Fab, there are six hypervariable loops - three from the Hc and three from the Lc - collectively referred to as complementarity-determining regions (CDRs) (22–24). The remaining domains of the Hc form the Fc (fragment crystallizable) region, which exhibits a higher degree of sequence conservation (22– 24). The antibody chains are assembled from four gene segments: Variable (V), Diversity (D, Hc only), Joining (J), and Constant (C). These segments undergo V(D)J recombination and somatic hypermutation, processes that generate extensive sequence diversity, particularly in the CDRs (21– 23). Among the three CDRs, CDR3 is the most diverse due to its length and positioning at the junction of the V, (D), and J gene segments (21–23). Given their central role in defining antibody specificity and enabling direct antigen recognition, accurate determination of CDR sequences is crucial (24, 25).

While sequencing a single Hc and Lc theoretically suffices to reconstruct an antibody, in practice this is hindered by biological and technical limitations (21, 23, 26). A key to successful proteomics-based antibody sequencing is obtaining enough redundant peptides covering the full sequence of the hypervariable CDR regions and their connection to constant regions. Standard proteolytic approaches, such as tryptic digestion, often fail to effectively fully cover these segments due to the limited sequence coverage, peptide length and redundancy (16, 25–29). A current way around this issue relies on the parallel use of multiple proteases for digestion, typically employing 4–6 proteases with complementary specificities. This multi-protease approach enhances sequence coverage by generating a broader array of overlapping peptides, thereby improving CDR resolution and overall sequence assignment (16, 26–28). However, it increases experimental and computational demands, reducing throughput and efficiency. Another challenge is the analysis of larger peptides (>1,500 Da), which are more commonly produced by non-tryptic proteases and, while analytically challenging, their greater sequence span is highly valuable for *de novo* sequencing. These longer peptides are more difficult to ionize and fully fragment using conventional collision-induced dissociation (CID/HCD), which reduces spectral quality and leads to gaps in the peptide sequence coverage (10, 26, 30, 31). To address these limitations, alternative fragmentation methods such as electron-based dissociation (ExD) have been shown to improve fragmentation efficiency for long, modified, and/or non-tryptic peptides (10, 26, 32–34). CID and ExD fragmentation schemes are complementary techniques that generate different fragment ion series across the peptide backbone, where CID produces b- and y-ions, while ExD yields c- and z-ions, providing orthogonal sequence information (10, 32–35). Building on this complementarity, hybrid fragmentation approaches, such as EThcD and EAciD, that combine ExD with supplemental collisional activation to produce rich, complementary spectra, have been introduced (34, 36, 37).

Despite these clear benefits, multi-protease and hybrid-fragmentation based strategies often require costly reagents, complex protocols, dedicated computing software, and specialized instrumentation, which can limit their accessibility and scalability (16, 27, 29). To address these limitations, we aimed here to develop a simplified strategy for *de novo* antibody sequencing, exploring two broadly specific hyperthermoacidic archaeal (HTA)-proteases, named Krakatoa and Vesuvius (38, 39) in combination with the ZenoTOF 7600 system MS. HTA-proteases are highly divergent from conventional proteases in both form and function, operating under extreme conditions that denature mesophilic proteins and enable rapid, one-step digestion without chaotropic agents and alkylation (38–40). Their broad specificity leads to long, diverse peptides ideal for *de novo* analysis. The ZenoTOF 7600 system enables tunable electron-activated dissociation (EAD) with short reaction times, which addresses the common drawbacks of longer duty cycles and lower scan rates associated with electron-transfer dissociation (34, 35). Moreover, EAD on the ZenoTOF 7600 system can be combined with supplemental CID activation, making it well-suited to obtain full sequences of even larger peptides (34, 35). As proof of concept, we optimized and applied this workflow to a mixture of four recombinant monoclonal antibodies (mAbs), as a mimic of an endogenous polyclonal mixture. Our HTA workflow generated highly redundant peptide reads, also across all variable CDR regions, enabling full sequence identification across four discrete antibodies in the mixture.

## Results and Discussion

Here, we explored the potential of two distinct HTA-proteases combined with hybrid fragmentation strategies on a ZenoTOF 7600 system for MS-based *de novo* antibody sequencing. The ultimate goal of this approach is to *de novo* sequence polyclonal antibody mixtures from bodily fluids, such as blood and saliva (15). Yet, additionally to the above-mentioned challenges, the complex nature of bodily fluids-derived samples hinders correct antibody sequence assignment, as digestion of polyclonal mixtures leads to a signal dilution of peptides from unique CDR regions. These peptides are overshadowed by the abundant peptides from the constant regions shared among nearly all antibodies, which substantially hampers the analysis. To mimic a simple polyclonal mixture, we mixed four mAbs of known sequence (Table S1): Cetuximab (CTX), Trastuzimab (TZB), NISTmAb, and F59. The first two are clinically used mAbs, the NISTmAb is a reference standard developed by the National Institute of Standards and Technology (NIST), and lastly, F59 is an in-house produced recombinant mAb, which we previously reported on (28).

We performed proteolytic digestion of the four mAbs mixture with the HTA-proteases, Krakatoa and Vesuvius. We benchmarked the performance of the HTA-proteases against trypsin and chymotrypsin, since the latter two are frequently used in proteomics and represent central components of multi-protease *de novo* sequencing approaches (25–27). As we aimed to generate long, highly overlapping peptides (38), we also explored MS methods to aid confident sequence assignment. To do so, we optimized hybrid fragmentation approach combining CID and EAD on a ZenoTOF 7600 system. We hypothesized that the hybrid EAciD method would yield dense, information-rich MS2 fragmentation spectra, thereby enhancing confident identification of the generated peptides (33, 35).

### Hybrid EAciD fragmentation increases identification rates especially for longer peptides

To evaluate and compare the performance of different fragmentation strategies on the ZenoTOF 7600 system, we analyzed the proteolytic digests of the 4 mAbs mixture by using CID, EAD, and hybrid EAciD fragmentation schemes (Figure 1, Table S2). Importantly, the MS methods were optimized prior to the experiment as described in Figure S1 and reported previously (33). While EAD slightly outperformed CID in MS2 to peptide spectrum matches (PSMs) rate, more CID PSMs passed the score filtering criteria (Byonic score ≥ 150 and LogProb ≥ 3), resulting in a similar performance of both CID and EAD methods (Figure 1A). The hybrid EAciD method surpassed both CID and EAD methods in the number of identified and passing PSMs. The higher success rate of EAciD identifications reflects the information-rich fragmentation spectra (Figure 1B), which were obtained without sacrificing the total number of sequenced peptides (Figure 1A, Table S1). As expected, CID yielded primarily b- and y-ions, and the ion coverage dropped for longer peptides (Figure 1B, S2). On the other hand, EAD resulted primarily in the c- and z-ions. Advantageously, the hybrid EAciD method provided mixed fragmentation spectra with similarly abundant b-, y-, c-, and z-ions (Figure 1B, S2). This feature resulted in higher confidence in EAciD peptide identifications, as indicated by higher success rates (Figure 1A). The added value of EAciD fragmentation is also evident from an illustrative comparison of the peptide’s MS2 spectra by different fragmentation methods (Figure 1C), indicating that each amino acid position was supported by multiple fragment ions in EAciD. Taken together, our results suggest that EAciD fragmentation on the ZenoTOF 7600 system provides better quality MS2 scans than solely EAD and CID, without sacrificing the number of fragmented peptides and duty cycle length (Figure 1A, Table S1). Therefore, we selected the optimized EAciD method for the subsequent analyses described below.

**Figure 1.**
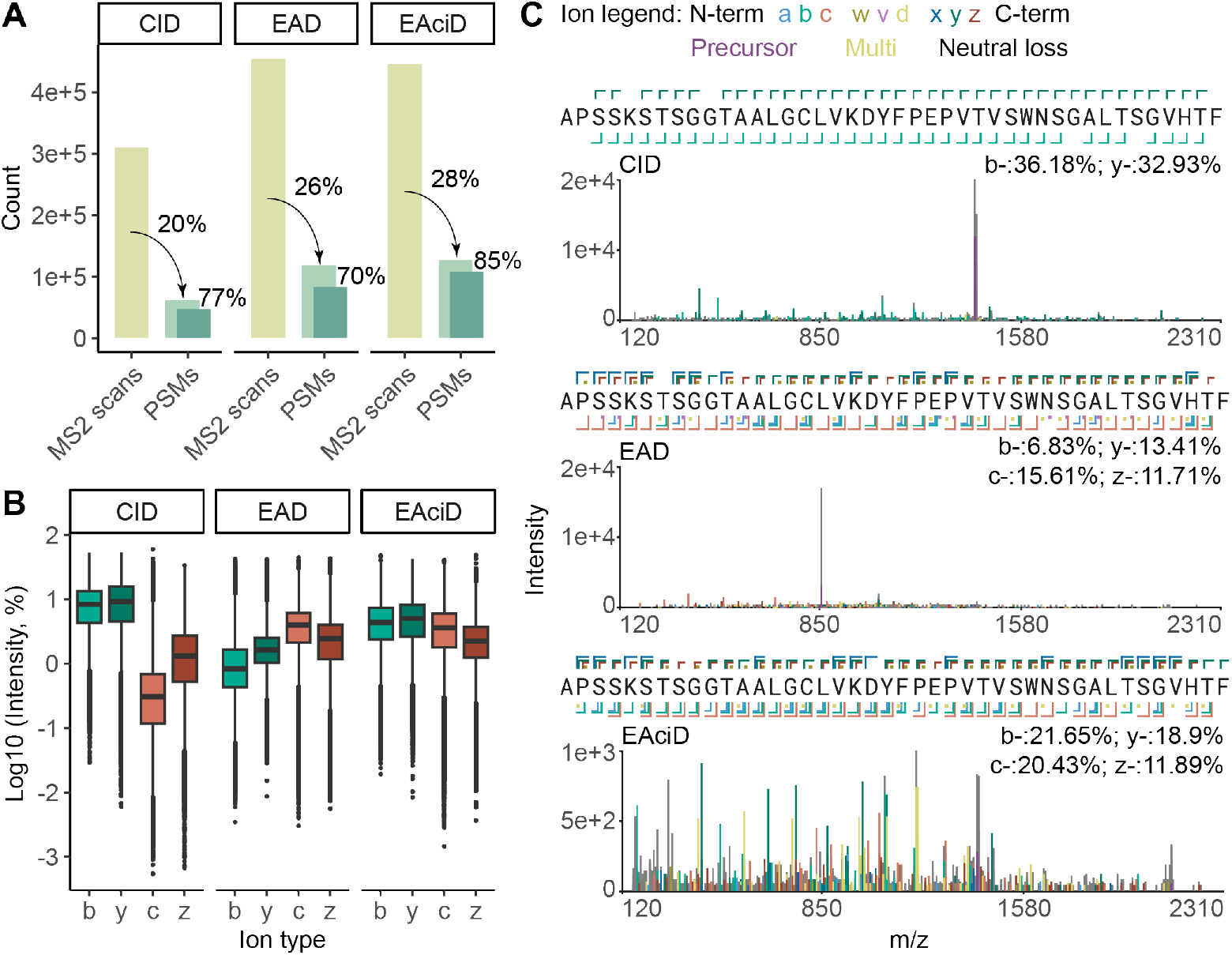
Evaluation of the fragmentation methods, CID, EAD and hybrid EAciD. Data shown in (A) and (B) are cumulative data acquired with the four used proteases on the sample mixture of the four antibodies. (A) MS2 scans and PSMs for each MS method. The percentages indicate the success rate of the identified MS2 scans (PSMs, light green), as well as the PSMs that meet the cutoff scores selected to filter the Byonic data (PSMs, dark green); Byonic score ≥ 150 and LogProb ≥ 3. EAciD has a higher success rate at both the MS2 scan and the PSM level, outperforming the single fragmentation schemes. (B) Intensity (%) of b-, c-, y-, and z-ions as observed in each of the fragmentation methods used. EAciD covers optimally all different b-, c-, y- and z-ion types. (C) Representative EAciD fragmentation spectrum of a 42 amino acid long peptide, originating from the TZB heavy chain, obtained through digestion with Krakatoa. The cartoon at the top of (C) shows the amino acid sequence, whereby always multiple fragment ions (e.g. b-, c-, y-, z-) cover each amino acid throughout the entire peptide.

### Single HTA-protease approach provides deep coverage with numerous unique and overlapping sequence reads

Our initial results clearly demonstrated the benefits of hybrid EAciD fragmentation, notably for longer peptides. These longer peptides are valued in *de novo* antibody sequencing since they may contain critical information to reconstruct CDR3 and assist with sequence assembly across more distant sequence variations in complex mixtures (24, 26, 27). Here, we assessed whether the broadly specific HTA-proteases Krakatoa and Vesuvius can generate such peptides and benchmarked the performance of the HTA-proteases against trypsin and chymotrypsin. We first evaluated the results by database matching search in Byonic, which is a conventional (non *de novo*) approach, providing high quality quantitative data and facilitating the comparison of the proteases.

We first assessed the total number of PSMs (Figure 2A, Table S2) obtained using the hybrid EAciD LC-MS method. In the tryptic digest, we detected a median of 5728 PSMs. The number of PSMs more than doubled for chymotrypsin and each of the HTA-proteases, with 13429, 11005, and 12236 PSMs, respectively for chymotrypsin, Vesuvius, and Krakatoa. This contrast likely reflects differences in protease cleavage preferences, with trypsin cleaving exclusively C-terminal of K and R, while the HTA-proteases exhibit broader cleavage specificities (Figure S3). Despite the high number of PSMs detected in the chymotrypsin sample, it resulted in only a slightly higher number of unique peptides compared to trypsin, namely 339 and 420 for trypsin and chymotrypsin, respectively (Figure 2A, S4, Table S2). In sharp contrast, both HTA-proteases produced up to five times more unique peptides than trypsin and chymotrypsin: 1675 and 2118 for Vesuvius and Krakatoa, respectively. These HTA peptides also cover a substantially wider range in both peptide lengths and charges (Figure 2B, S5-S6), with the median peptide length (26 amino acids) exceeding that observed when using trypsin (18 amino acids) or chymotrypsin (20 amino acids) by 44% and 30%, respectively.

**Figure 2.**
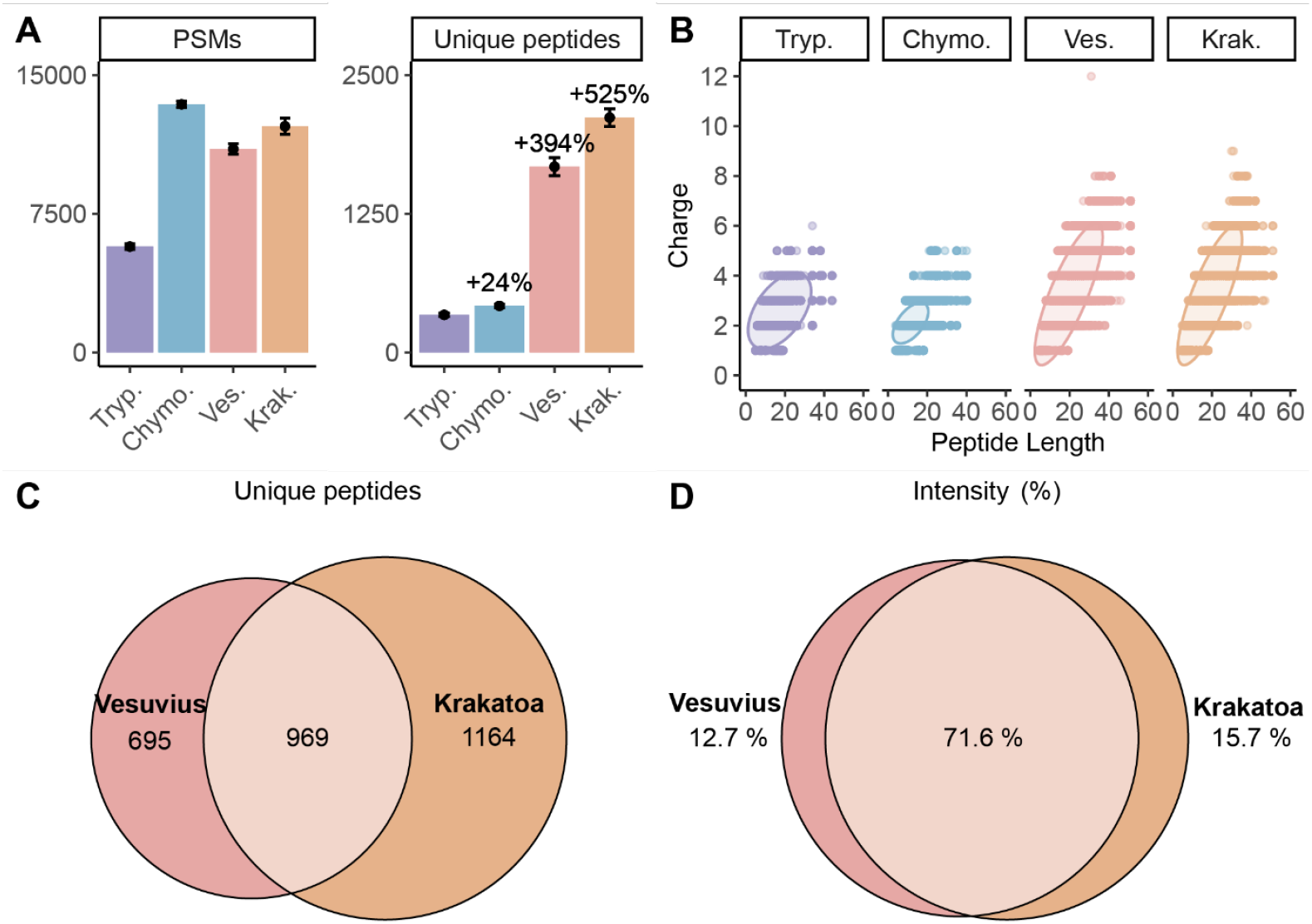
Peptide data for the mAb mixture following parallel digestion with the 4 proteases: trypsin (purple), chymotrypsin (blue), Vesuvius (pink), and Krakatoa (orange). (A) Obtained peptide spectrum matches (PSMs) (left) and unique peptides (right) identified per protease. The percentages represent increases in unique peptides, when compared to trypsin. (B) All unique peptides were analyzed based on their charge and peptide length. Digestion with Vesuvius and Krakatoa provides substantially broader distributions of peptides both in the peptide length and charge dimension. The ellipses represent the 0.95 confidence level of a multivariate t-distribution. (C) Venn diagram representing the overlap and unique peptides identified by using Vesuvius and Krakatoa. (D) The overlapping peptides generated by both Vesuvius and Krakatoa are generally the more abundant ones. This data originates solely from the data generated in EAciD mode, filtered for Byonic score ≥150 and log ≥ 3 from n= 3 technical replicates.

As previously reported (37) and observed in our data (Figure S3), Krakatoa and Vesuvius exhibit similar cleavage preferences at the C-terminal of E, F, L, D, M, and Q. Moreover, both proteases yielded a similar number of PSMs and unique peptides, resulting in comparable peptide distributions (Figure 2A-B). When we assessed the overlap of the HTA-generated peptides (Figure 2C-D), we observed that only 34% of unique peptides (n = 969) were shared among Krakatoa and Vesuvius, although they account for 72% of MS1 intensity. This indicates that each HTA-protease generates a slightly different profile of peptides, with Krakatoa marginally outperforming Vesuvius in both PSMs and the number of unique peptides. From the 4-5 fold increase in unique peptide sequence reads obtained by HTA-proteases, we proposed that the multi-protease approach typically used in *de novo* antibody sequencing (24, 26, 27) might be effectively replaced by using a single HTA-protease.

To further assess the suitability of HTA-proteases for *de novo* sequencing, we analyzed the redundancy of unique peptide reads covering the antibody sequences (Figure 3, S7). We adopted the redundancy metric, as it is the key parameter for successful *de novo* sequence assembly (15, 26). In this context, each additional unique read of the same sequence enhances the confidence and increases the accuracy of the assignment. Figure 3 visualizes the number of redundant reads obtained from unique peptides generated by the HTA-proteases, trypsin, or chymotrypsin. Strikingly, both HTA-proteases provided numerous unique overlapping peptide-reads confidently covering the crucial CDRs in addition to the rest of the antibody framework. Of note, we observed slightly higher redundancy for the constant regions shared among multiple antibodies in our sample, when compared to unique variable CDRs for all proteases. Consequently, neither trypsin, chymotrypsin, nor their combination covered all CDR regions (Figure 3). Yet, each HTA-protease covered all CDR regions of all antibodies with several unique reads. Notably, the confident and redundant coverage of CDR regions, which is of particular interest for *de novo* sequencing, was achieved even with a single HTA-protease for all four mixed mAbs (Figure S7-S8).

**Figure 3.**
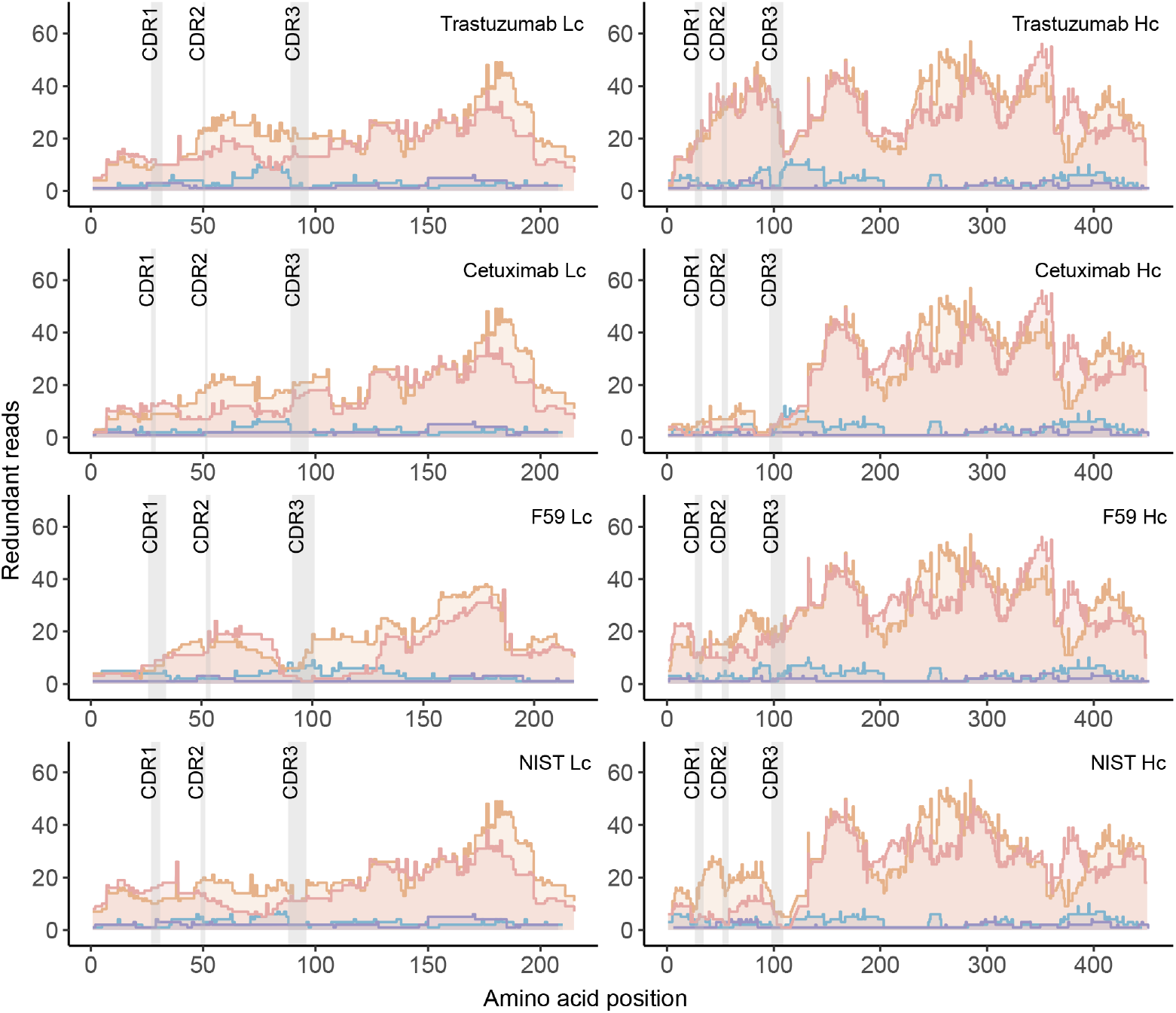
Redundant reads per amino acid. Each graph depicts the number of times each amino acid in the sequence of one of the antibody chains is covered by unique peptides. The data displayed originates from the digest by trypsin (purple), chymotrypsin (blue), Vesuvius (pink), or Krakatoa (orange). The HTA-proteases, Vesuvius and Krakatoa, consistently provide substantially more sequence reads for all the light and heavy chains of the studied antibodies, including the CDR regions. The individual reads, including both shorter and longer reads, are depicted.

The superior performance of HTA-proteases is also evident from a global redundancy analysis of all 4 mAbs in the sample (Figure 4A). While trypsin and chymotrypsin yielded only a median of 1 and 3 unique redundant reads per amino acid (redundancy of 1 and 3), respectively, the median redundancies obtained by HTA-proteases were 23 and 24 for Vesuvius and Krakatoa, respectively. Notably, the minimum redundancy observed in the Krakatoa sample was 2, indicating that this single HTA-protease covered all CDR regions as well as the entire sequence of all 4 mAbs with at least 2 unique peptides covering each amino acid in the full sequence.

**Figure 4.**
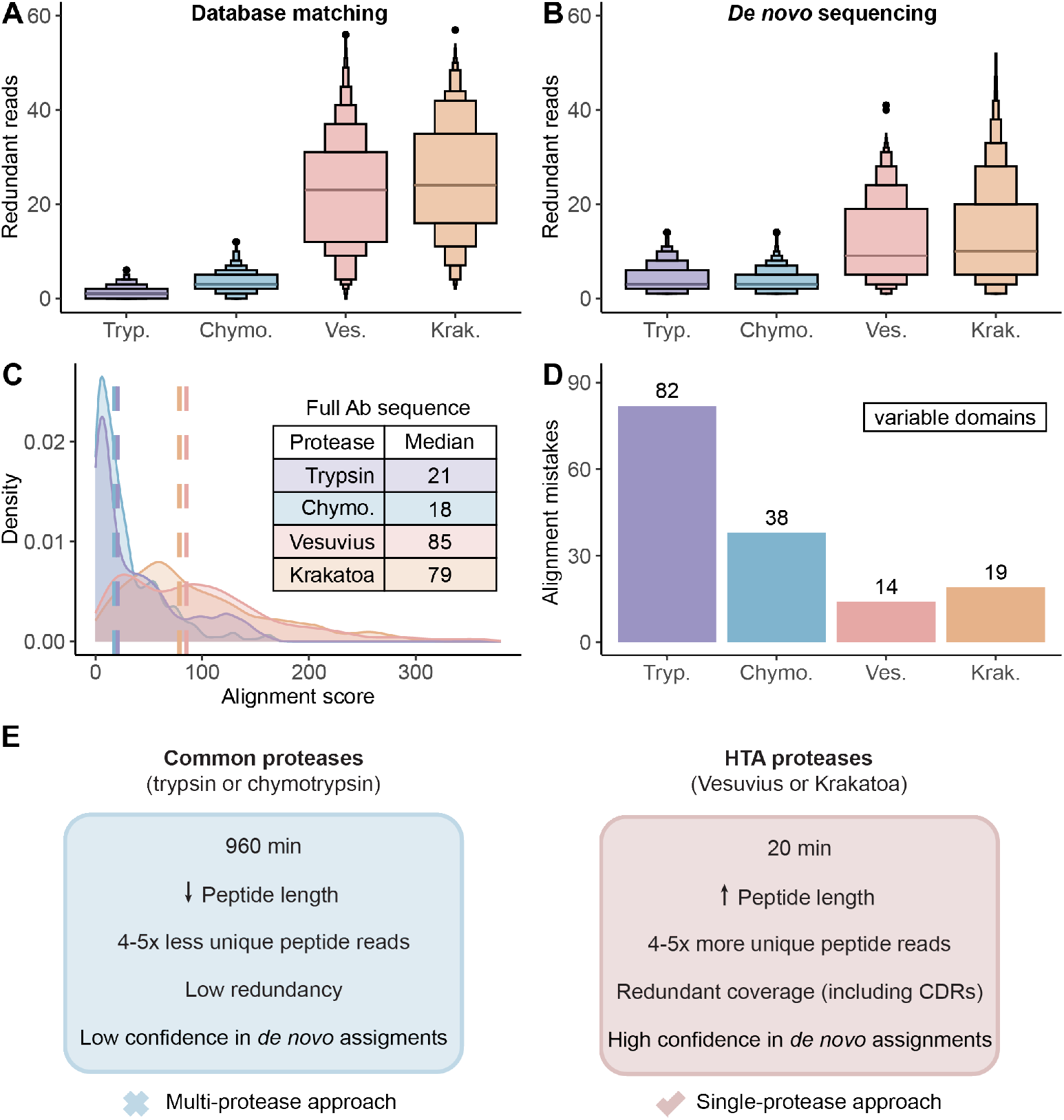
HTA–proteases combined with EAciD provide redundant and confident *de novo* reads. (A) Distribution of the unique redundant reads per protease, cumulative data from analyzing both the Lc and Hc chains of all four investigated antibodies. The median number of redundant reads corresponds to 1, 3, 23, and 24 for trypsin, chymotrypsin, Vesuvius, and Krakatoa, respectively. (B) Distribution of the unique *de novo* reads per protease, cumulative data from both the Lc and Hc chains of all four investigated antibodies. Unique d*e novo* reads were defined based on the amino acid position on the sequences; only peptides identical to the antibody consensus sequence were included. The median number of redundant reads corresponds to 3, 3, 9, and 10 for trypsin, chymotrypsin, Vesuvius, and Krakatoa, respectively. Unique *de novo* reads for the complete *de novo* dataset shows comparable results as indicated in Figure S10. (C) Kernel density distribution of the maximum alignment scores for each amino acid as provided by Stitch. The scores are calculated as the sum of the positional score for all reads in the alignment. (D) Sum of the alignment mistakes detected when matching the template to the consensus sequence, for the variable regions of all four antibodies. (D) Summary of the characteristics of the proteases used for digesting the mAb mixture. The table shows the time required for the proteases to achieve sufficient protein digestion and the number of reads obtained.

### HTA-protease approach enhances confidence in *de novo* antibody sequencing

Seeing the superior performance of HTA-proteases in the database matching search, we sought to perform template-independent *de novo* sequencing on our dataset. We used PEAKS with the DeepNovo/De Novo algorithm (40) followed by *de novo* reconstruction of the antibody sequences in Stitch (41). Of note, most of the currently available *de novo* peptide search algorithms, including PEAKS, have been tailored for CID/HCD fragmented tryptic peptides (42) hampering the data analysis for alternative approaches. Still, we pursued the *de novo* analysis, mainly focusing on antibody variable regions for their critical role in antigen binding and consequently, of particular importance in *de novo* sequencing.

To evaluate the suitability of our HTA-protease/EAciD approach for *de novo* sequencing, we assessed several crucial metrics as redundancy, alignment scores, and alignment mistakes (Figure 4). All these factors together determine the suitability of a given workflow for the successful *de novo* sequence assembly. Consistent with the database matching results (Figure 4A-B, S9-S10), the median number of redundant reads increased more than threefold for the HTA-proteases, compared to trypsin and chymotrypsin. A similar trend was also observed in the alignment score, where the HTA-proteases achieved four times higher scores than trypsin and chymotrypsin (Figure 4C). This metric represents the confidence of each amino acid assignment in the four mAb sequences, showing that the HTA-proteases provide less ambiguity in the final assembled sequences. The higher confidence of sequence assignment using the HTA-based approach was also reflected in the alignment mistakes (Figure 4D). Here, we assessed errors produced in the amino acid assignment of the template sequence compared to the consensus for all variable regions of the four mAbs. Remarkably, the number of mistakes reduced up to fourfold when using the HTA-proteases combined with EAciD, showing that the high redundancy and variety of HTA-generated peptides enable more accurate *de novo* sequence assignments with fewer errors.

In summary, our results show that the HTA-protease/EAciD approach provides extensive redundancy, CDRs coverage accompanied by higher alignment scores and less alignment mistakes. These results, obtained by using a single HTA-protease, are comparable to those typically generated using multiple conventional enzymes in *de novo* sequencing (26). Furthermore, our HTA approach offers additional benefits: very short digestion times, no need for extensive sample cleanup, and analysis just by a single EAciD LC-MS/MS run (Figure 4E). An additional advantage of the HTA-based workflow is that the samples are handled at low pH, keeping cysteines protonated. Therefore, cysteine alkylation is not only unnecessary but also chemically unfeasible, thereby simplifying the workflow and facilitating data analysis. These properties make the HTA-based approaches highly time-, cost-, and sample-efficient, representing a significant advancement for *de novo* sequencing applications, even beyond antibody analysis.

## Conclusions

The vast majority of MS-based bottom-up proteomics currently relies on database-matching approaches, which are dependent on previously identified DNA-based sequences. This is a limiting factor for analyzing proteins without an easily accessible DNA-template such as alternative splicing products, genetic variants, endogenously processed proteins, and importantly humoral antibody repertoires. All these proteins require a *de novo* sequencing approach, where the sequence is directly determined from the MS2 data. MS-based *de novo* sequencing of antibodies is currently feasible, especially when targeting a single purified mAb, but remains a cost- and labor-intensive challenge. It requires unambiguous and confident identifications of numerous unique and overlapping peptides that enable the *de novo* assembly of a complete sequence. To obtain enough sequence reads, an antibody sample is typically digested in parallel by several proteases, and respective digests are independently analyzed by LC-MS/MS.

Here, we benchmarked a novel approach to simplify antibody *de novo* sequencing. We used the HTA-proteases Krakatoa and Vesuvius to generate redundant peptide reads and implemented a hybrid EAciD peptide fragmentation method to facilitate peptide identification in a single LC-MS/MS run. Our results demonstrated that both HTA-proteases generate a diverse array of peptide reads, exceeding the redundancies achieved by conventionally used proteases. By combining the optimized EAciD method with HTA-proteases, we detected 4-5 times more unique peptides than with trypsin and chymotrypsin. These unique HTA-protease reads covered all CDR regions as well as the entire sequences of all four mAbs in one sample with high redundancy both in the database and *de novo* searches. Moreover, we observed a fourfold increase in *de novo* alignment scores, accompanied by a similar decrease in the alignment mistakes in the HTA samples compared to trypsin. Our results represent a significant improvement in redundancy and sequence coverage, compared to trypsin and chymotrypsin, indicating that a single HTA-protease can provide superior performance to conventional multi-protease approaches. While we focused solely on antibody *de novo* sequencing, the use of HTA-proteases should be equally advantageous across a broad range of bottom-up proteomics applications.

## Methods

### Proteases and antibodies used

The HTA-proteases, Krakatoa and Vesuvius, were provided by CinderBio and have been introduced previously (37, 38). These enzymes are recombinantly produced and purified to >98% based on protease activity. Trypsin was acquired from Promega and chymotrypsin from Sigma. F59 mAb was recombinantly produced by Genmab, Utrecht, as described earlier (28) based on a sequence retrieved by *de novo* sequencing. Trastuzumab (TZB) was provided by Roche. Cetuximab (CTX) was acquired from Evidentic and NISTmAb was purchased from Agilent. The sequences of all four mAbs are provided in Table S1. No unexpected or unusually high safety hazards were encountered.

### Antibody digest preparation

As a simplified model for an endogenous polyclonal mixture, four mAbs of known sequence: trastuzumab (TZB), NISTmAb, cetuximab (CTX), and F59 (28) were mixed in equimolar ratio. For the HTA digestion, this antibody mixture was diluted into the reducing HTA digestion buffer (1x vendor-provided HTA acidic buffer, 5 mM TCEP, 2 Units/1 µg of protein of Krakatoa protease or 10 Units/1 µg of protein of Vesuvius protease), followed by incubation at 80 °C for 20 min, shaking at 350 rpm. The digestion was stopped by snap-freezing in liquid nitrogen. For the trypsin and chymotrypsin digestion, the same antibody mixture was denatured, reduced and alkylated and subjected to a SP3 digestion according to a published protocol (43). Trypsin or chymotrypsin was added at 1:50 ratio (protease: antibody, w/w) and incubated overnight at 37 °C, shaking at 1000 rpm. Peptides were acidified with 0.2% TFA (final concentration), collected and frozen in liquid nitrogen until further use.

### LC-MS/MS analysis

The generated peptides (100 ng) were analyzed using a ZenoTOF 7600 system (SCIEX) mass spectrometer coupled to an Ultimate3000 UHPLC system (ThermoFisher). The peptides were concentrated and desalted by using a PepMap™ Neo Trap Cartridge (5 mm × 0.3 mm, 5 μm, Thermo Fisher Scientific) trap column, followed by their separation using an IonOpticks Aurora Elite XS C18 analytical column (15 cm x 75 µm) with a 65-min gradient, starting at 3% buffer B (0.1% formic acid in 80% acetonitrile) to 9% B in 2 min, from 9% to 30% in 37 min, from 30% to 44% in 15 min, from 44% to 55% in 4 min, from 55% to 99% in 1 min, 99% wash-out in 5 min and re-equilibration back to 3% buffer B in 10 min where buffer A is 0.1% formic acid in water. The flow rate was 300 nl/min. The ZenoTOF 7600 system nanoflow source conditions were as follows: spray voltage, 1500 V; nano gas 1, 10 psi; curtain gas, 35 psi; CAD gas, 7; nano cell temperature, 300 °C; column temperature, 30 °C. The parameters for the MS1 scans were as follows: mass range, *m/z* 350-3000; accumulation time, 0.05 s; declustering potential, 80 V; collisional energy (CE), 10 V; time bins to sum, 8. MS2 spectra were acquired with CID, EAD, and EAciD fragmentation schemes. For the EAD and EAciD, the 15 most intense precursors with a minimum intensity of 50 counts per second (cps) and with charge states between 2-10 were selected for fragmentation. For CID, 25 monoisotopic top candidates with a minimum intensity of 300 cps with the same charge states (2-10) were selected for fragmentation. The MS2 parameters for all fragmentation methods were as follows: Zeno trapping, On; Zeno threshold, 100000 cps; TOF mass range, *m/z* 150-3000; exclusion for 6 s after 1 occurrence. MS2 parameters for EAD and EAciD were as follows: electron beam current, 7000 nA; electron kinetic energy (KE), 9 eV; reaction time, 20 ms; accumulation time, 65 ms. For the CID fragmentation scheme, accumulation time was set to 0.02 s. For the CID and EAciD schemes, supplemental collisional energy was set to default dynamic CE for MS/MS.

### MS data analysis

The generated wiff2 files were searched in Byonic (Protein Metrics, v5.7.11). A focused database containing all four Hc and Lc of TZB, NISTmAb, CTX, F59, and keratins was used in the search. The cleavage specificity was set to semi-specific C-terminally of L/F/D/M/E/Q/I for Krakatoa and Vesuvius, allowing 6 missed cleavages. For trypsin, the cleavage was set to fully specific C-terminally of K/R, allowing 2 missed cleavages. For chymotrypsin, the cleavage was set to C-terminally of F/L/M/I/Y/W, allowing 6 missed cleavages. Carbamidomethyl was set as a fixed modification for trypsin and chymotrypsin; variable modifications for all the proteases were as follows: M and W oxidation as common 2; pyroGlu on N-terminus of protein for E and Q as common 1; deamidation of N as rare 1; N-glycan (57 human plasma N-glycan database) as rare 1. A total of 3 common and 2 rare modifications were allowed per peptide. The mass tolerance was set to 20 ppm for both MS1 and MS2 scans. Fragmentation was set as EThcD for the EAciD files, ECD for the EAD files and QTOF/HCD for the CID files. The peptide identifications were accepted with a score ≥ 150 and LogProb ≥ 3. The peak areas for the peptide abundance analysis were retrieved in Skyline (v. 24.1.0.414) (44). MS2 Spectrum annotation was performed by using multi-annotator (45) to retrieve ion type information on all fragment ions. For this purpose, wiff files were converted to mzML format using the MSConvert tool from ProteoWizard (46), with the peakPicking filter enabled.

For *de novo* search, the wiff files were searched in Peaks 11 software (Bioinformatics Solutions Inc., Waterloo, ON, Canada) using PEAKS DeepNovo/De Novo algorithm. The search was performed using unespecific enzyme for the HTA – Proteases; fragmentation method EAD for the EAciD files; precursor mass 10ppm; fragment ion 0.02 Da; Oxidation (HW), oxidation (M), pyro-glu (Q, E) were set as variable modifications; allowing a maximum of 4 PTMs per peptide. Carbamidomethylation (C) was set as a fixed modification for trypsin and chymotrypsin. Data was exported from PEAKS, filtered for an ALC score of 80 and run in the in-house built software tool Stitch in order to reconstruct the antibody sequences from the *de novo* peptides (41). The filter parameters used in Stitch were Cutoffscore 8 and EnforceUnique 0.8 for Template Matching. The Stitch source code and a detailed manual are accessible via GitHub (https://github.com/snijderlab/stitch).

The data analysis and visualization were performed with custom R- and python scripts. Data were analyzed in R version 4.4.0 running in RStudio 2024.12.1 (Build 563). Data handling and visualization were mostly performed with *tidyverse* (v2.0.0) and *cowplot* (v1.1.3). The venn diagram and the boxplots were made with the packages *eulerr* (v7.0.2) and *lvplot* (v0.2.1), respectively. In Figure 3B the ellipses represent the 0.95 confidence level of a multivariate t-distribution, based on the *stat_ellipse* function from *ggplot2* in R.

## Data availability

LC-MS/MS data have been deposited to the ProteomeXchange Consortium via the PRIDE partner repository (47) with the dataset identifier PXD063988.

## Supporting Information

The supporting information file contains additional Figures S1-S9 and Tables S1-S2, as referenced in the text.

## Declaration of possible conflicts of interests

P. Pribil and S. Heidelberger are employees of Sciex, the manufacturer of the ZenoTOF 7600 system used in this study; S.M. Yannone is the inventor of HTA-technologies and serving as CEO at CinderBio, and Allison Michele Narlock-Brand is employed at CinderBio, the company that produces and supplies the HTA-proteases commercially. The remaining authors declare no competing interests.

## Acknowledgements

We would like to thank Shelley Jager (Utrecht University, NL) for helpful suggestions regarding the data acquisition and analysis. The recombinant F59 used in this study was recombinantly expressed, purified, and kindly provided by Aran Labrijn and Boris Bleijlevens (Genmab, Utrecht, NL). We thank Dietmar Reusch and Markus Haberger (Roche, Penzberg, DE) for the kind donation of trastuzumab. The graphical abstract was created in BioRender (Shamorkina, T. (2025) https://BioRender.com/hgpe1k6). This research received funding by the Netherlands Organization for Scientific Research (NWO) through the Spinoza Award SPI.2017.028 to AJRH, Gravitation 2013 BOO, Institute for Chemical Immunology (ICI; 024.002.009) to JS, and the European Union through an ERC-2023-ADG grant REVAMP nr: 101141457, to AJRH, and by the National Institute of General Medical Sciences of the National Institutes of Health under award number 5R44GM128540 (SMY).

## Graphical abstract

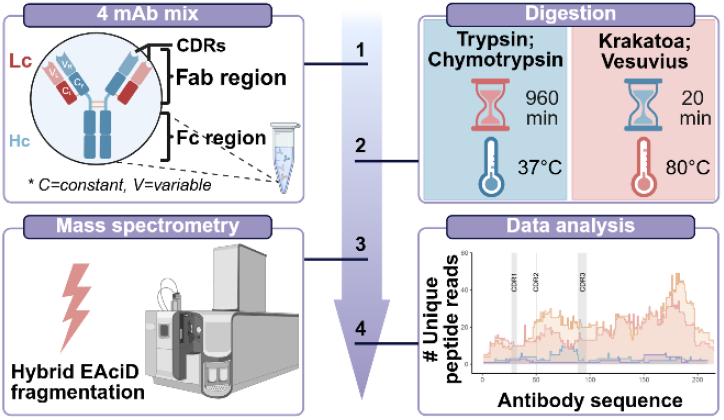

### Synopsis

Efficient hyperthermal acidic proteases provide five times more unique peptide reads than trypsin or chymotrypsin, greatly advancing *de novo* antibody sequencing.

