## supplementary information for "Deep Coverage and Extended Sequence Reads Obtained with a Single Archaeal Protease Expedite *de novo* Protein Sequencing by Mass Spectrometry"


### Contents

#### Figures


S1. Optimization of kinetic energy (KE) parameters for the EAciD method
S2. EAciD performs especially well on long-read peptides
S3. Sequence logos for (A) Krakatoa and (B) Vesuvius protease cleavage specificities
S4. Representative LC-MS chromatograms of the peptide digest of the four-antibody mixture
S5. Peptide length distribution per protease
S6. Peptide charge distribution per protease
S7. Redundancy in sequence coverage by unique peptide reads
S8. CDR regions coverage by HTA proteases
S9. Redundant reads per amino acid as obtained by true *de novo* sequencing
S10. Unique *de novo* reads per protease


#### Tables


S1. Sequences of the heavy and light chains of the antibodies used in the four monoclonal antibody mixture
S2. Summary of unique peptides, PSMs, and MS2 scans observed in each combination of protease and MS fragmentation method

### Supplementary Figures

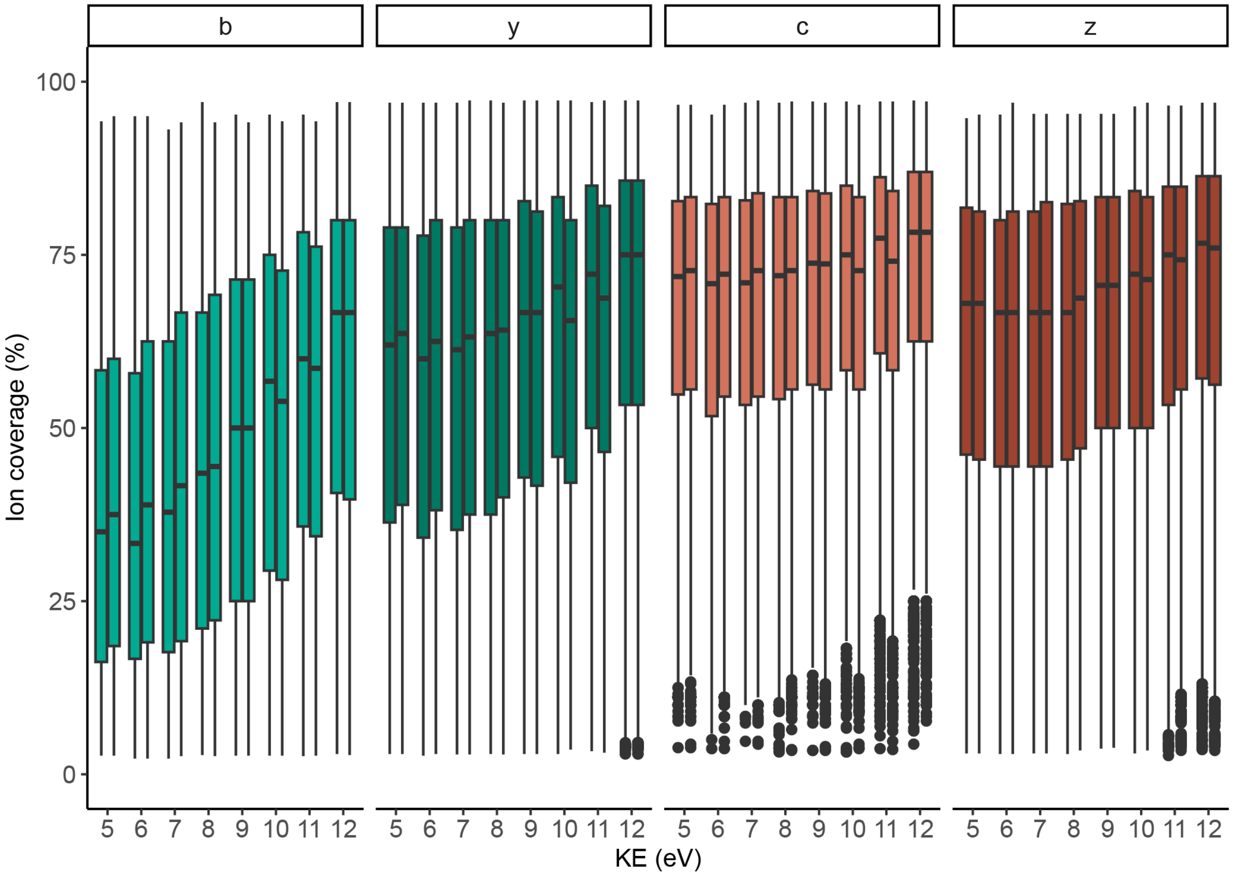


**Figure S1. Optimization of kinetic energy (KE) parameters for the EAciD method.** The figure depicts the median peptide sequence coverages (y-axis) by ion series (b-, y-, z-, and c-ions) originating from a Krakatoa digest. The data were collected during optimization of the EAD method using various kinetic energies ranging from 5 to 12 eV (x-axes). As kinetic energy increases, the proportion of b-ions in the EAD fragmentation also rises, indicating enhanced secondary fragmentation and greater occurrence of neutral losses. We chose 9 eV as an intermediate kinetic energy setting to complement the supplemental collisional activation for the hybrid EAciD approach. The data originates from two replicate runs acquired at each kinetic energy setting.

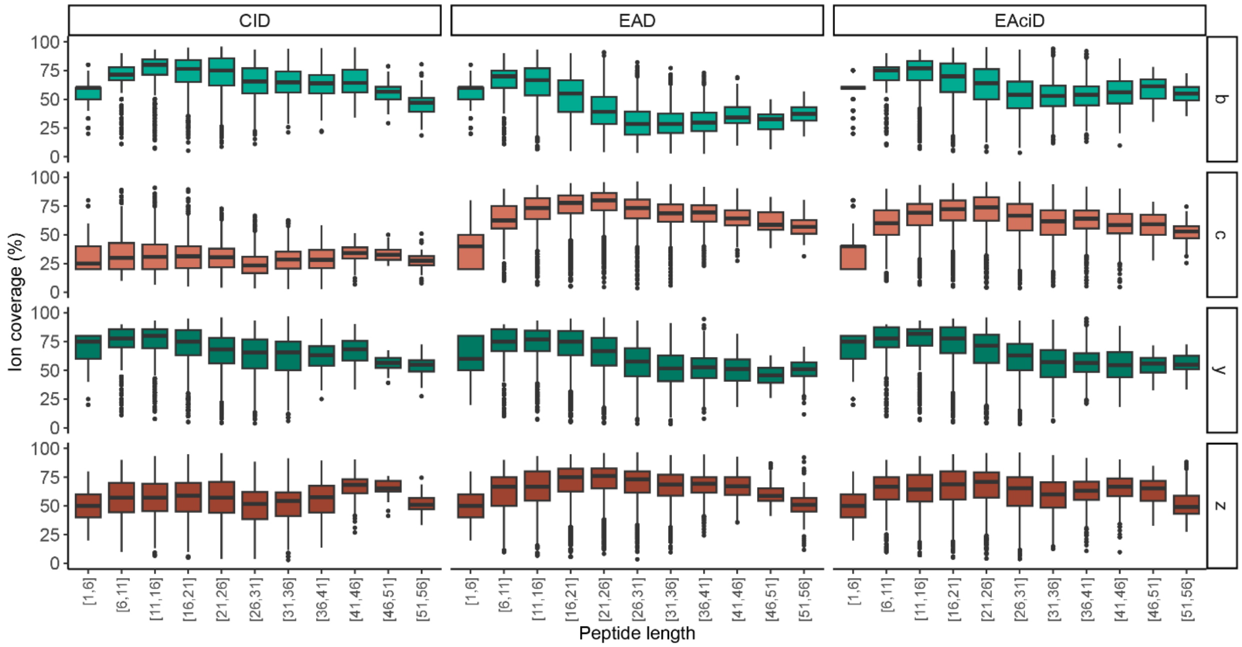


**Figure S2. EAciD performs especially well on long peptide reads**. Sequence coverage across different peptide ion lengths for the fragmentation methods tested. Results show that CID yielded primarily b- and y- ion series, while c- and z- ion series are prevalent in EAD. The hybrid use of these two fragmentation methods, EAciD, provides good ion coverage for all b-, y-, c- and z- ion series. Of note, the presence of b- and y-ions in pure EAD fragmentation and c-and z-ions in pure CID fragmentation indicates the degree of secondary fragmentation and neutral losses (1–3).

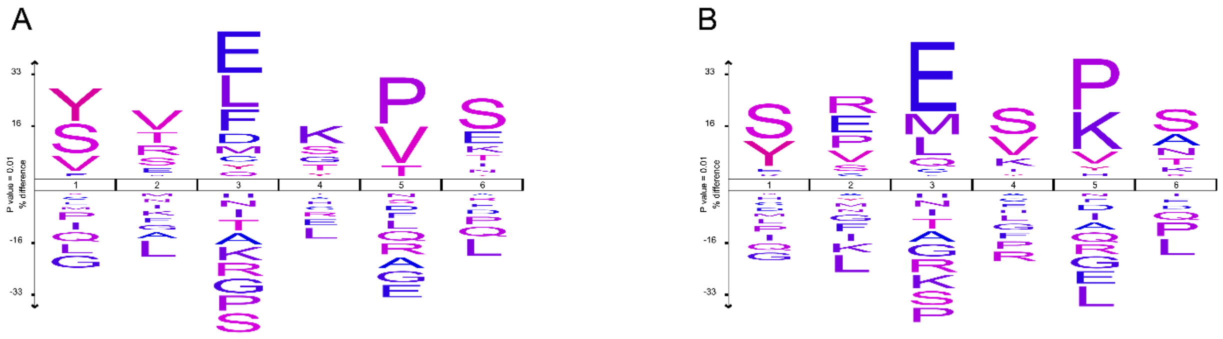

**Figure S3. Sequence logos for (A) Krakatoa and (B) Vesuvius protease cleavage specificities**. The specificities were determined for the data generated from the four mAb mixture analyzed by EAciD method. The logos were generated with IceLogo (4) against the human reference proteome with a 0.01 P-value cutoff. The cleavage site is localized C-terminal of the 3rd position. The observed cleavage specificities are similar to previously published data (5, 6). The minor differences in the cleavage specificities, when compared to previous reports, arise most likely from using here a simple sample consisting of just four alike mAbs.

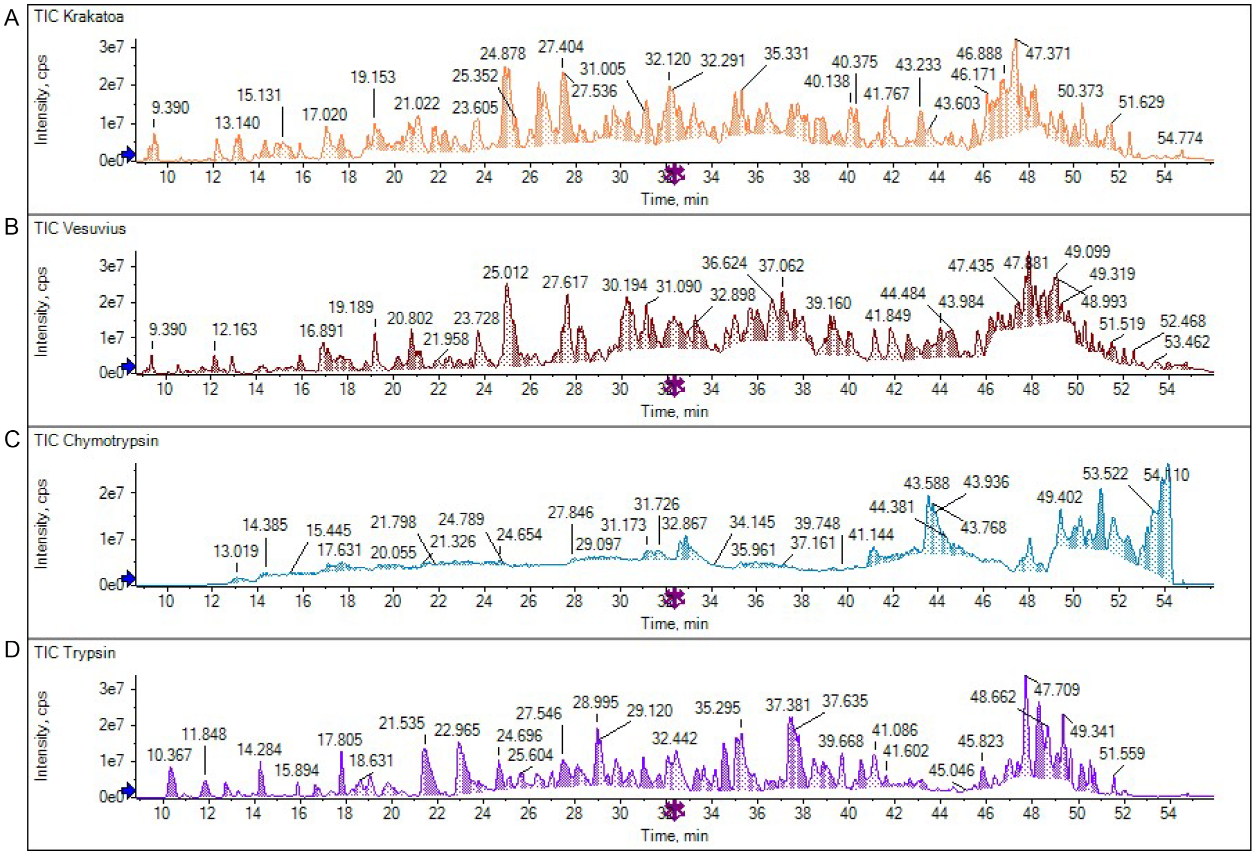


**Figure S4. Representative LC-MS chromatograms of the peptide digest of the four-antibody mixture.** LC-MS chromatograms of the Krakatoa (orange) and Vesuvius (dark red) digests display a high density of well-resolved peaks, surpassing the overall peak complexity observed in the tryptic digest (purple). In contrast, the LC-MS chromatograms of chymotrypsin (blue) were consistently of equal abundance but revealed less peaks, that generally showed more tailing. Importantly, all LC-MS conditions (settings and buffers used) were identical in all runs.

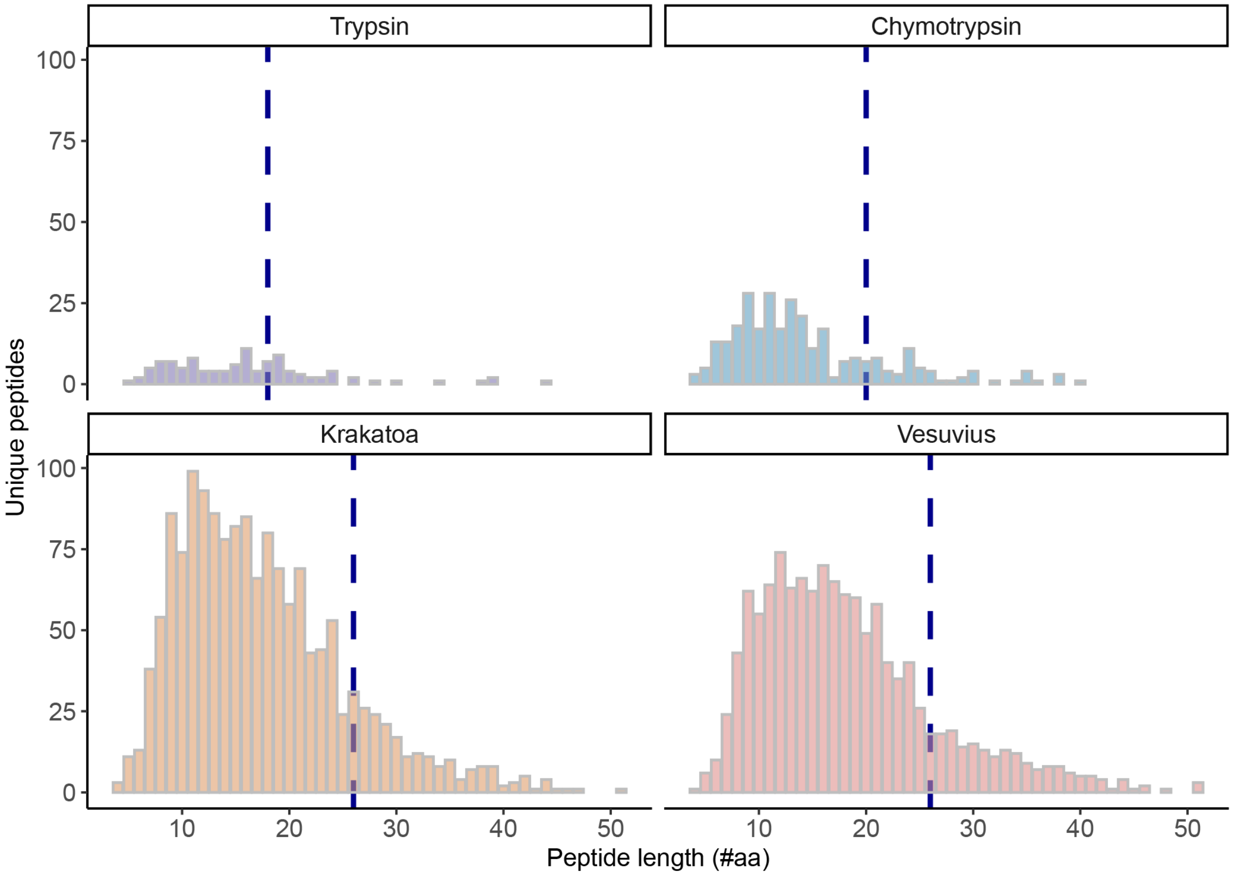


**Figure S5. Peptide length distribution per protease.** The bar plot depicts the peptide length distribution of the unique peptides identified per protease. Vesuvius and Krakatoa show a higher number of unique peptide identifications and a broader peptide length distribution, compared to trypsin and chymotrypsin. Remarkably, the HTA-proteases also produce a substantial number of longer peptides, resulting in a higher median peptide length, which is of great value in *de novo* sequencing to achieve better sequence coverage. The median ranges are 18, 20, 26 and 26 respectively for trypsin, chymotrypsin, Vesuvius, and Krakatoa, as indicated by the blue line. This data originates solely from the data generated in EAciD mode, filtered for Byonic score ≥150 and log ≥ 3 from n= 3 technical replicates.

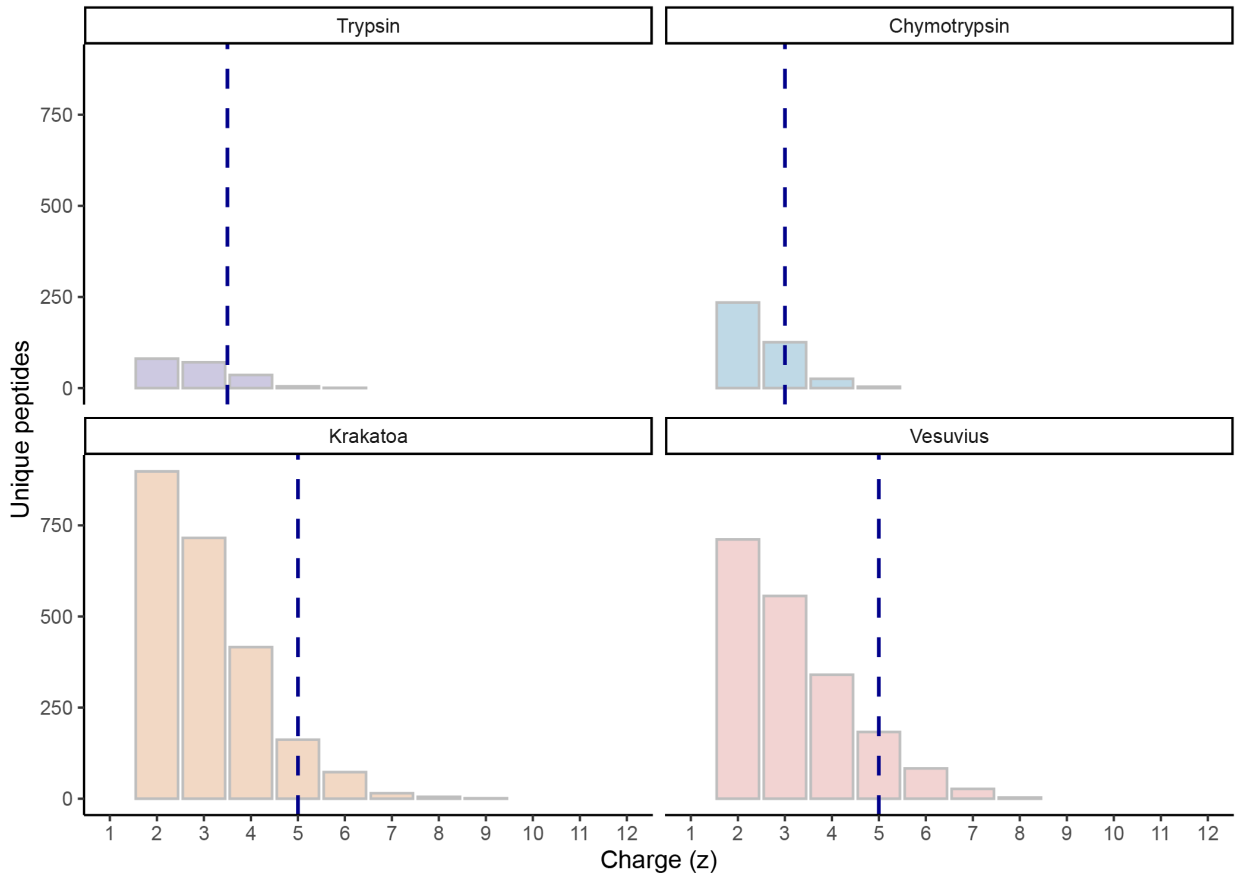


**Figure S6. Peptide charge distribution per protease.** The bar plot depicts the charge distribution of the unique peptides identified per protease. Vesuvius and Krakatoa show a higher number of unique peptide identifications and a considerable broader charge distribution, compared to trypsin and chymotrypsin. The median ranges are +3.5, +3.0, +5.0 and +5.0 respectively for trypsin, chymotrypsin, Vesuvius, and Krakatoa; as indicated by the blue line. This data originates solely from the data generated in EAciD mode, filtered for Byonic score ≥150 and log ≥ 3 from n= 3 technical replicates.

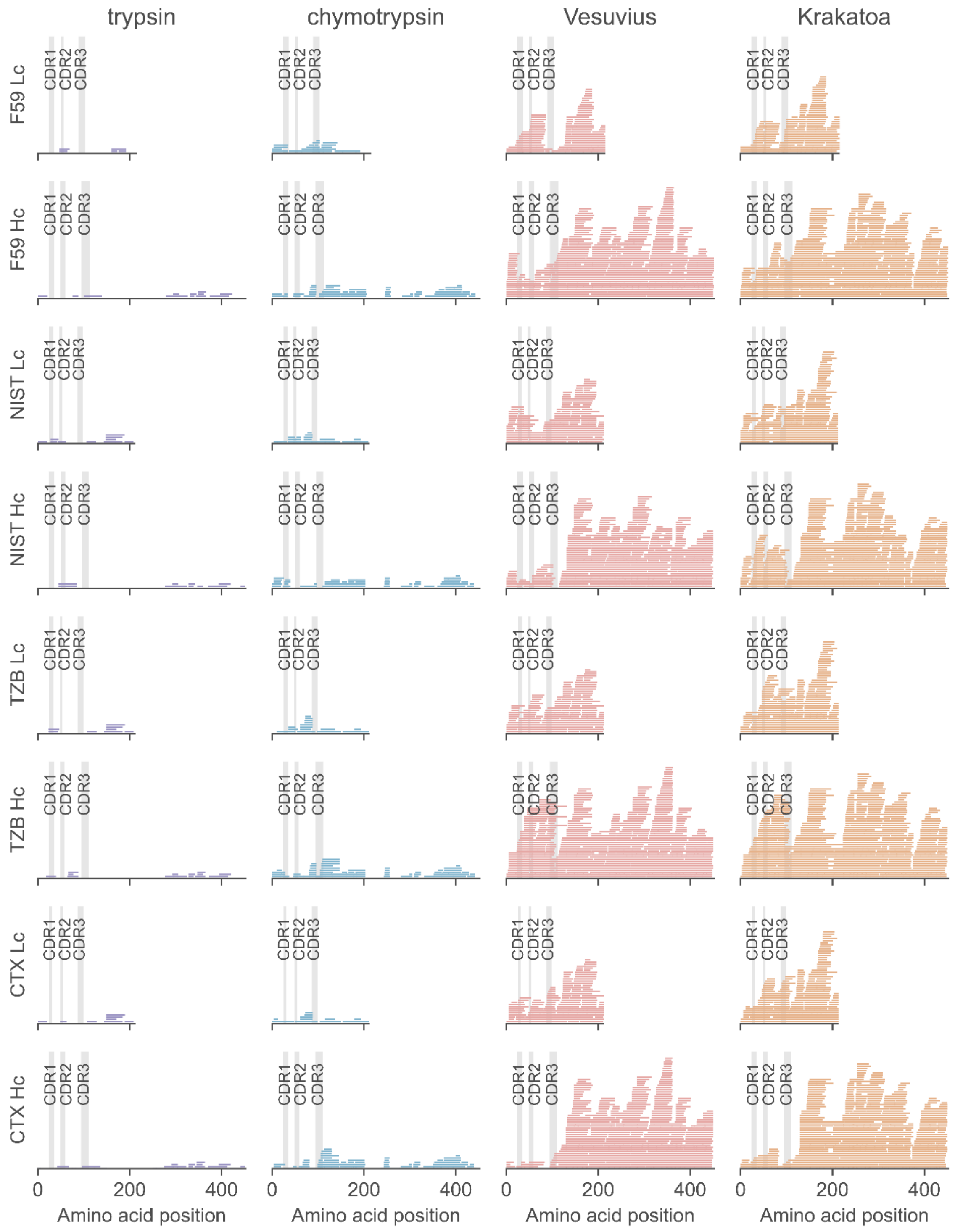


**Figure S7. Redundancy in sequence coverage by unique peptide reads**. Depicted are the unique peptides generated by trypsin (purple), chymotrypsin (blue), Vesuvius (red), and Krakatoa (orange) spanning over the Lc and Hc sequences of all four mAbs. Each unique peptide detected is visualized as a line covering the respective region in the protein sequence. CDR regions are highlighted in gray and labeled. This data originates solely from the data generated in EAciD mode, filtered for Byonic score ≥150 and log ≥ 3 from n= 3 technical replicates.

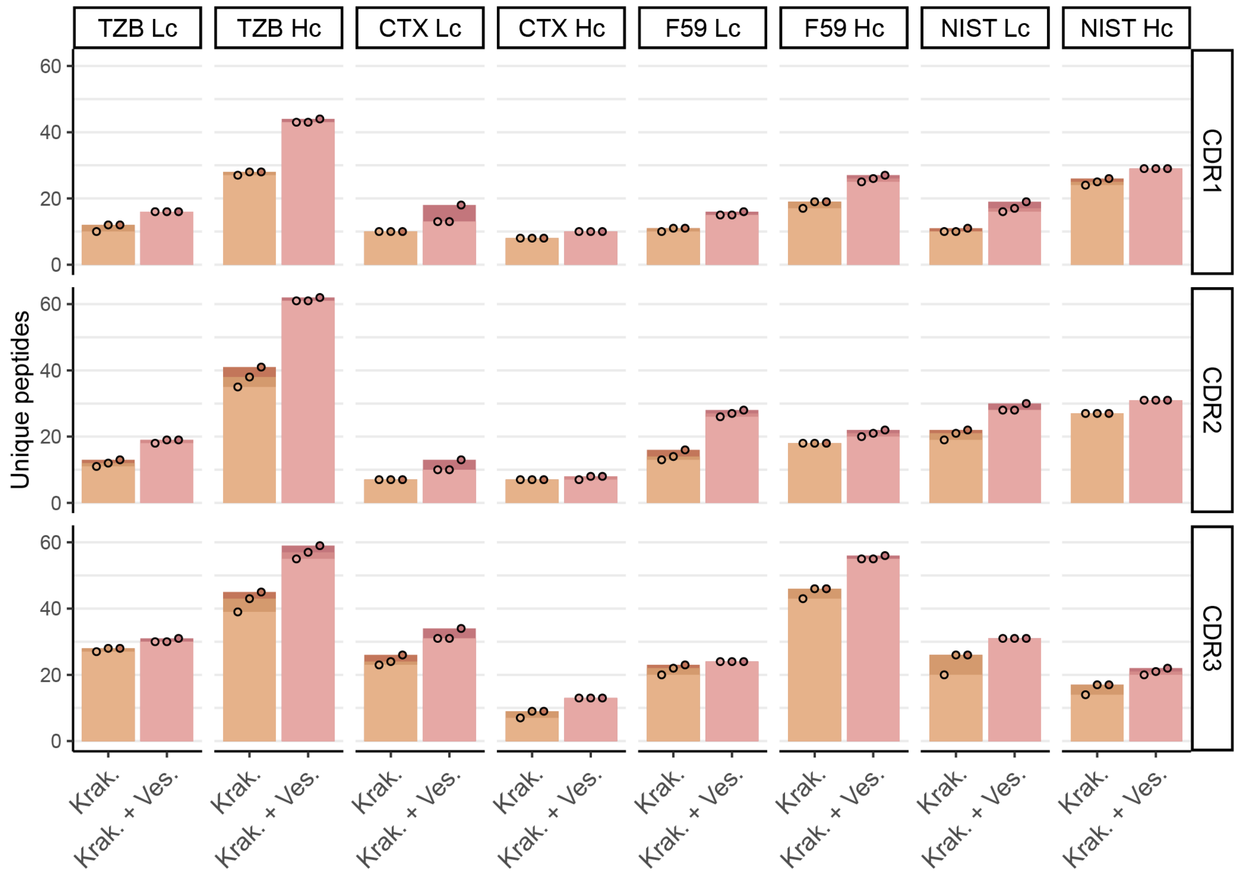


**Figure S8. CDR regions coverage by HTA proteases.** The sum of unique peptides is shown covering the CDR regions per individual antibody, either by using Krakatoa or by combining data from the Krakatoa and Vesuvius digest. Although the combined usage of the two proteases marginally increases the number of identified unique peptides, data from a single Krakatoa digest are sufficient to fully cover all CDR regions. Each black circle represents the cumulative unique peptides detected per replicate. This data originates solely from the data generated in EAciD mode, filtered for Byonic score ≥150 and log ≥ 3 from n= 3 technical replicates.

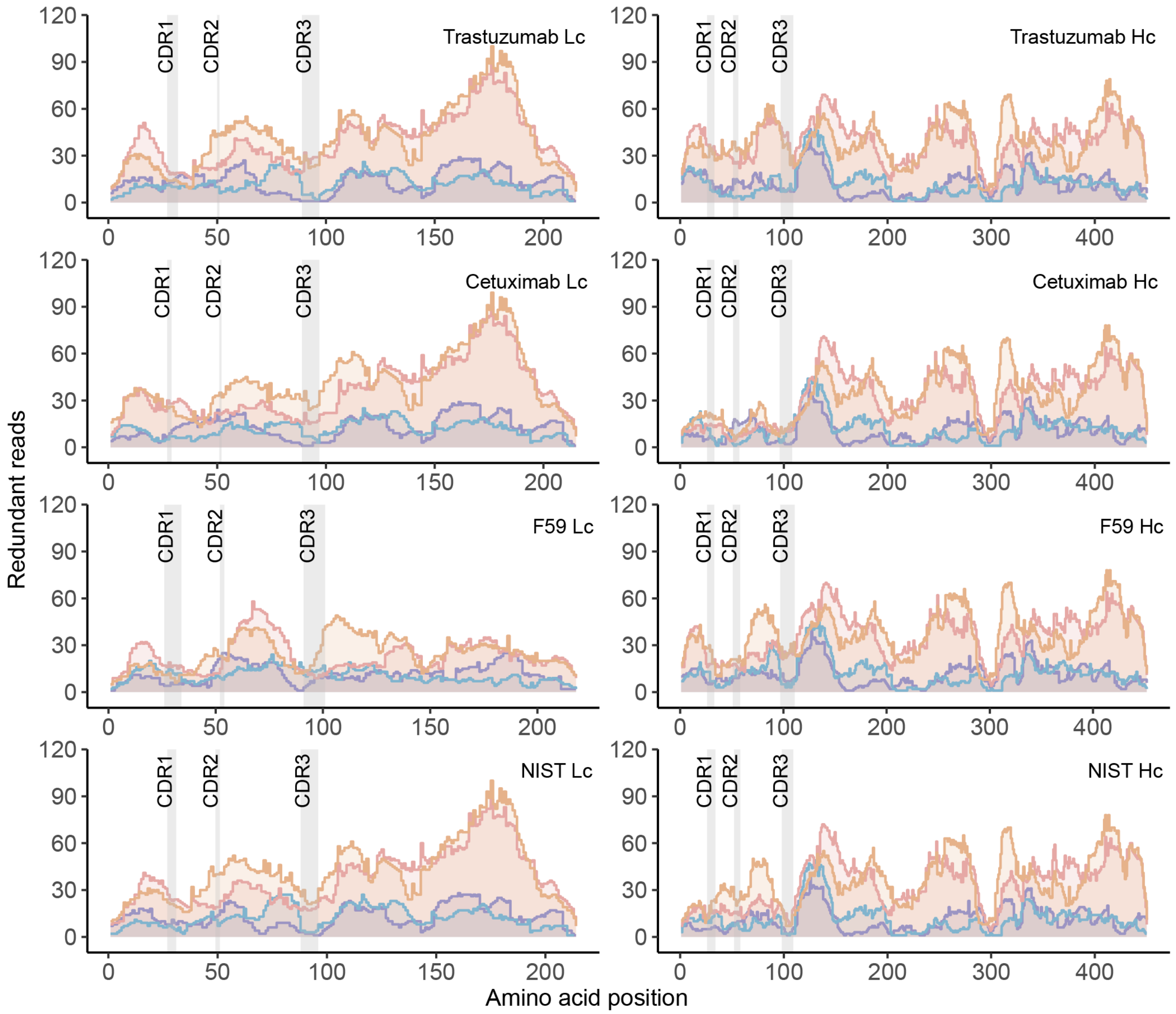


**Figure S9. Redundant reads per amino acid as obtained by true de novo sequencing.** Each graph depicts the number of times each amino acid in the sequence of one of the antibody chains is covered by unique de novo annotated peptides. The data displayed originates from the digest by trypsin (purple), chymotrypsin (blue), Vesuvius (pink), or Krakatoa (orange). The HTA-proteases, Vesuvius and Krakatoa, consistently provide substantially more *de novo* sequence reads for all the light and heavy chains of the studied antibodies, including the CDR regions. The individual reads, including both shorter and longer reads, are depicted. This data originates solely from the data generated in EAciD mode, filtered for PEAKS ALC score ≥80, and for Stitch Cutoff Score 8 and Enforce Unique 0.8 for Template Matching, from n= 3 technical replicates.

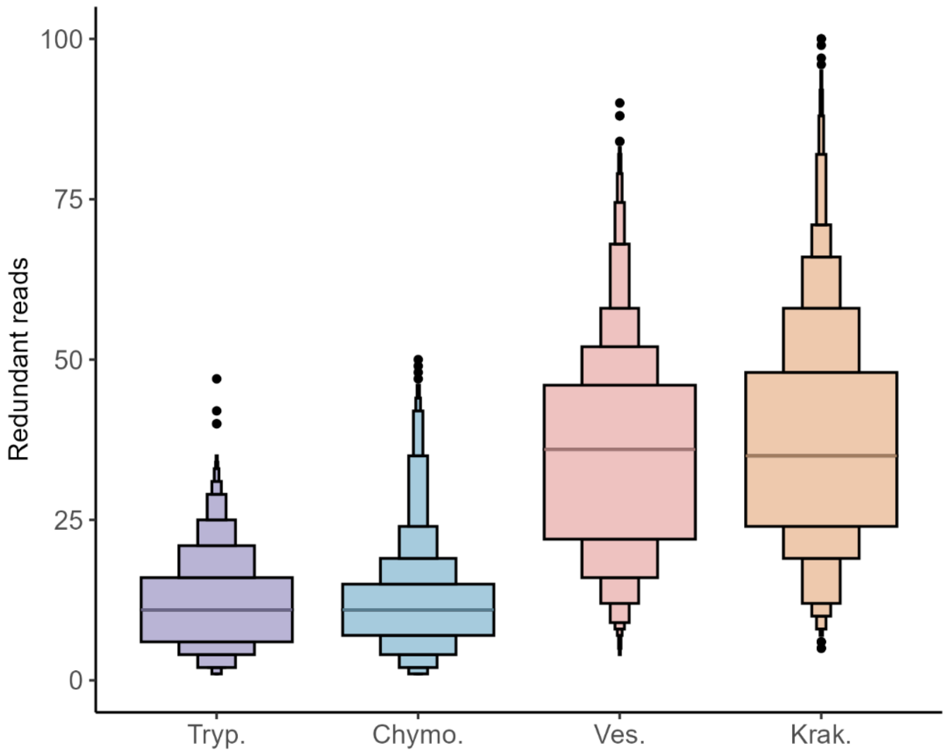


**Figure S10. Unique *de novo* reads per protease.** Cumulative data from both the Lc and Hc chains of all four investigated antibodies. *De novo* unique reads were defined based on the amino acid position on the sequences; all peptides that, based on Stitch annotation, did not start with insertions or deletions compared to the consensus sequence were included. The median number of redundant reads corresponds to 11, 11, 36, and 35 for trypsin, chymotrypsin, Vesuvius, and Krakatoa, respectively.

### Supplementary Tables

**Table S1. Sequences of the heavy and light chains of the antibodies used in the four monoclonal antibody mixture.**

| Description | Sequence |
|---|---|
| Trastuzumab (Light chain) | DIQMTQSPSSLSASVGDRVTITCRASQDVNTAVAWYQQKPGKAPKLLIYSASFLYSGVPSRFSGSRSGTDFTL TISSLQPEDFATYYCQQHYTTPPTFGQGTKVEIKRTVAAPSVFIFPPSDEQLKSGTASVVCLLNNFYPREAKVQ WKVDNALQSGNSQESVTEQDSKDSTYSLSSTLTLSKADYEKHKVYACEVTHQGLSSPVTKSFNRGEC |
| Trastuzumab (Heavy chain) | EVQLVESGGGLVQPGGSLRLSCAASGFNIKDTYIHWVRQAPGKGLEWVARIYPTNGYTRYADSVKGRFTISA DTSKNTAYLQMNSLRAEDTAVYYCSRWGGDGFYAMDYWGQGTLVTVSSASTKGPSVFPLAPSSKSTSGGT AALGCLVKDYFPEPVTVSWNSGALTSGVHTFPAVLQSSGLYSLSSVVTVPSSSLGTQTYICNVNHKPSNTKVD KKVEPKSCDKTHTCPPCPAPELLGGPSVFLFPPKPKDTLMISRTPEVTCVVVDVSHEDPEVKFNWYVDGVEV HNAKTKPREEQYNSTYRVVSVLTVLHQDWLNGKEYKCKVSNKALPAPIEKTISKAKGQPREPQVYTLPPSREE MTKNQVSLTCLVKGFYPSDIAVEWESNGQPENNYKTTPPVLDSDGSFFLYSKLTVDKSRWQQGNVFSCSVM HEALHNHYTQKSLSLSPG |
| Cetuximab (Light chain) | DILLTQSPVILSVSPGERVSFSCRASQSIGTNIHWYQQRTNGSPRLLIKYASESISGIPSRFSGSGSGTDFTLSINS VESEDIADYYCQQNNNWPTTFGAGTKLELKRTVAAPSVFIFPPSDEQLKSGTASVVCLLNNFYPREAKVQWK VDNALQSGNSQESVTEQDSKDSTYSLSSTLTLSKADYEKHKVYACEVTHQGLSSPVTKSFNRGEC |
| Cetuximab (Heavy chain) | QVQLKQSGPGLVQPSQSLSITCTVSGFSLTNYGVHWVRQSPGKGLEWLGVIWSGGNTDYNTPFTSRLSINK DNSKSQVFFKMNSLQSNDTAIYYCARALTYYDYEFAYWGQGTLVTVSAASTKGPSVFPLAPSSKSTSGGTAAL GCLVKDYFPEPVTVSWNSGALTSGVHTFPAVLQSSGLYSLSSVVTVPSSSLGTQTYICNVNHKPSNTKVDKRV EPKSCDKTHTCPPCPAPELLGGPSVFLFPPKPKDTLMISRTPEVTCVVVDVSHEDPEVKFNWYVDGVEVHNA KTKPREEQYNSTYRVVSVLTVLHQDWLNGKEYKCKVSNKALPAPIEKTISKAKGQPREPQVYTLPPSREEMTK NQVSLTCLVKGFYPSDIAVEWESNGQPENNYKTTPPVLDSDGSFFLYSKLTVDKSRWQQGNVFSCSVMHEA LHNHYTQKSLSLSPGK |
| F59 (Light chain) | QSALTQPASVSGSPGQSITISCTGTSSDVGGYNYVSWYQHHPGKAPKLLISEVSDRPSGVSSRFSGSKSGNTA SLTISGLQAEDESMYFCSSYTDLTSFVVFGGGTKLTVLGQPKAAPSVTLFPPSSEELQANKATLVCLISDFYPGA VTVAWKADSSPVKAGVETTTPSKQSNNKYAASSYLSLTPEQWKSHRSYSCQVTHEGSTVEKTVAPTECS |
| F59 (Heavy chain) | EPELVESGGGLAQPGTSLRLSCEASGFTFDDYAMHWVRQAPGRALEWVSGISWSSDNLAYSDSVEGRFTIS RDNAKNSLYLQMNSLRLDDTAFYYCAKDVRPYYDFWAFDSWGRGTPVTVSSASTKGPSVFPLAPSSKSTSG GTAALGCLVKDYFPEPVTVSWNSGALTSGVHTFPAVLQSSGLYSLSSVVTVPSSSLGTQTYICNVNHKPSNTK VDKRVEPKSCDKTHTCPPCPAPELLGGPSVFLFPPKPKDTLMISRTPEVTCVVVDVSHEDPEVKFNWYVDGV EVHNAKTKPREEQYNSTYRVVSVLTVLHQDWLNGKEYKCKVSNKALPAPIEKTISKAKGQPREPQVYTLPPSR EEMTKNQVSLTCLVKGFYPSDIAVEWESNGQPENNYKTTPPVLDSDGSFFLYSKLTVDKSRWQQGNVFSCS VMHEALHNHYTQKSLSLSPG |
| NIST (Light chain) | DIQMTQSPSTLSASVGDRVTITCSASSRVGYMHWYQQKPGKAPKLLIYDTSKLASGVPSRFSGSGSGTEFTLTI SSLQPDDFATYYCFQGSGYPFTFGGGTKVEIKRTVAAPSVFIFPPSDEQLKSGTASVVCLLNNFYPREAKVQW KVDNALQSGNSQESVTEQDSKDSTYSLSSTLTLSKADYEKHKVYACEVTHQGLSSPVTKSFNRGEC |
| NIST (Heavy chain) | QVTLRESGPALVKPTQTLTLTCTFSGFSLSTAGMSVGWIRQPPGKALEWLADIWWDDKKHYNPSLKDRLTIS KDTSKNQVVLKVTNMDPADTATYYCARDMIFNFYFDVWGQGTTVTVSSASTKGPSVFPLAPSSKSTSGGTA ALGCLVKDYFPEPVTVSWNSGALTSGVHTFPAVLQSSGLYSLSSVVTVPSSSLGTQTYICNVNHKPSNTKVDK RVEPKSCDKTHTCPPCPAPELLGGPSVFLFPPKPKDTLMISRTPEVTCVVVDVSHEDPEVKFNWYVDGVEVH NAKTKPREEQYNSTYRVVSVLTVLHQDWLNGKEYKCKVSNKALPAPIEKTISKAKGQPREPQVYTLPPSREMT KNQVSLTCLVKGFYPSDIAVEWESNGQPENNYKTTPPVLDSDGSFFLYSKLTVDKSRWQQGNVFSCSVMHE ALHNHYTQKSLSLSPGK |

**Table S2. Summary of unique peptides, PSMs, and MS2 scans observed in each combination of protease and MS fragmentation method.** Values represent the median across three replicates.

| Protease | MS method | Unique peptides | PSMs | MS2 scans |
|---|---|---|---|---|
| Trypsin | CID | 313 | 2264 | 16251 |
| Trypsin | EAD | 322 | 4712 | 32115 |
| Trypsin | EAciD | 339 | 5728 | 31765 |
| Chymotrypsin | CID | 581 | 8527 | 29182 |
| Chymotrypsin | EAD | 451 | 13661 | 40600 |
| Chymotrypsin | EAciD | 420 | 13429 | 40497 |
| Vesuvius | CID | 1777 | 6113 | 32826 |
| Vesuvius | EAD | 1541 | 9919 | 39401 |
| Vesuvius | EAciD | 1675 | 11005 | 38497 |
| Krakatoa | CID | 1246 | 3492 | 25956 |
| Krakatoa | EAD | 1943 | 11472 | 39725 |
| Krakatoa | EAciD | 2118 | 12236 | 38359 |
